# SSB1 links stress granule regulation to cellular stress responses and renal ischemia-reperfusion injury

**DOI:** 10.64898/2026.05.22.726875

**Authors:** János Pálinkás, Bálint Jezsó, Eszter Nagy-Kanta, Rita Németh, Usman Ashraf Awan, Gergely Takács, Bernadett Szikriszt, Ádám Tamás Hosszú, Péter Ecsédi, Gergely Szakács, Dávid Szüts, Andrea Fekete, Mihály Kovács

## Abstract

Mammalian single-stranded DNA binding protein 1 (SSB1) has been established as an essential component of genome stability in both human cells and mice. Moreover, SSB1 was recently implicated in cytoplasmic stress response by its involvement in Ras GTPase-activating protein-binding protein 1 (G3BP1)-containing cytoplasmic stress granules (SGs) upon various forms of stress. Here, we generated and analyzed human cellular knockout and rodent ischemia-reperfusion (I/R) models to define SSB1’s roles in cytoplasmic stress response. Analysis of wild-type as well as SSB1 and G3BP1 knockout human retinal pigment epithelial (RPE-1) cells shows stress-specific incorporation of SSB1 into SGs and a negative regulator role for SSB1 in SG dynamics under sublethal stress conditions. We find that SSB1 knockout measurably increases cellular sensitivity to oxidative stress but does not alter cell proliferation following mild acute stress. Moreover, we detect SSB1 efflux from the nucleus upon stress that is dependent upon the presence of G3BP1 in a stress-specific manner. In addition, using mouse and rat models we observe significant upregulation and robust cytoplasmic granulation of SSB1 upon renal ischemia-reperfusion stress, establishing SSB1’s involvement in complex organismal stress response *in vivo*. Together, our data demonstrate active involvement of SSB1 in cytoplasmic response to cellular stress and acute kidney injury, with implications for targeting stress response functions in cancerous *versus* non-cancerous contexts.

**HIGHLIGHTS:**

- SSB1 is incorporated into cytoplasmic stress granules and negatively regulates stress granule assembly under sublethal stress conditions
- SSB1 shows stress- and G3BP1-dependent nuclear efflux
- SSB1 is upregulated and undergoes apical granulation in renal epithelial cells during renal ischemia–reperfusion injury

## INTRODUCTION

Mammalian SSB1 (NABP2, OBFC2B, SOSS-S1) protein has been established as an essential factor supporting genome stability in human cell lines^1–9^ as well as knockout mouse models^10–15^. These studies have indicated SSB1’s involvement in homologous recombination (HR)-based repair of double-stranded DNA breaks (DSBs), base excision repair (BER) of oxidative DNA lesions as well as transcription termination control^3,16–20^. Notably, ablation-based assessment of SSB1’s cellular functions has mostly been limited to effects of siRNA-based SSB1 knockdown in cancer cell lines (HeLa, U2OS) on hydrogen peroxide (H_2_O_2_) and ionizing radiation survival^2,6,21–24^. Stable, clonal knockout models for SSB1 have been lacking, even though a previous study reported stress sensitization effects upon lentivirus/Cas9-based SSB1 ablation^6^.

Besides SSB1’s definitive roles in genome maintenance, our previous study provided evidence for the involvement of the protein in cytoplasmic stress response associated with stress granule (SG) formation in human cell lines^25^. In addition, we detected a redox-regulated liquid-liquid phase separation (LLPS) mechanism for SSB1 that could adaptively underlie cellular regulatory functions^25^. SGs are dynamic, membraneless cytoplasmic biomolecular condensates that assemble rapidly in response to a wide range of cellular stressors^26,27^. The resulting condensates sequester non-translating mRNAs alongside diverse RNA-binding proteins, translation initiation factors, and signaling molecules, enabling cells to prioritize survival-promoting gene expression programs^27^. At the biophysical level, SG formation is driven by LLPS, a ubiquitous process in cell biology, during which macromolecules undergo concentration-dependent demixing from the surrounding cytoplasm to form a distinct, condensed liquid-like phase^28,29^. LLPS is now recognized as a fundamental organizing principle underlying the formation of diverse membraneless organelles and its dysregulation is increasingly linked to diseases ranging from neurodegeneration to cancer^28,29^. SG assembly and disassembly are reversible: upon stress resolution, SGs rapidly dissolve and normal translation resumes, whereas failure to disassemble has been mechanistically linked to pathological protein aggregation observed in neurodegenerative diseases such as amyotrophic lateral sclerosis (ALS) and frontotemporal dementia (FTD)^30^.

A pivotal regulator of SG biogenesis is Ras GTPase-activating protein-binding protein 1 (G3BP1), a multidomain scaffold protein^31,32^. Upon cellular stress, binding of partner proteins or post-translational modifications enable G3BP1 dimerization and RNA-driven liquid-liquid phase separation (LLPS) to nucleate SGs ^31,32^. Consistently, cells lacking both G3BP1 and its paralog G3BP2 are largely deficient in forming canonical stress granules, firmly establishing G3BP proteins as essential SG nucleators^32,33^. Beyond their acute stress-adaptive functions, SGs and G3BP1 have been implicated in the regulation of mTOR signaling, apoptotic pathways, and oncogenesis, with G3BP1 found to be overexpressed in multiple cancer types^30^.

Our initial discoveries about SSB1 enrichment in SGs upon cellular stress motivated us in the current study to generate and utilize human cellular knockout systems as well as a mouse ischemia-reperfusion model to define the roles of SSB1 in cytoplasmic stress responses. Ischemia-reperfusion (I/R) injury is among the most common causes of acute kidney injury (AKI), a severe clinical syndrome characterized by rapid decline in renal function that carries high morbidity and mortality rates and for which specific therapeutic options remain limited^34–36^. Renal I/R inflicts complex, multiphasic cellular damage: the ischemic phase deprives tubular epithelial cells of oxygen and metabolic substrates, while the subsequent reperfusion paradoxically exacerbates injury through a burst of reactive oxygen species (ROS) production, mitochondrial dysfunction, inflammatory cytokine signaling, and the activation of regulated cell death pathways^36,37^. A hallmark of the renal I/R response is the concurrent engagement of both oxidative stress and ER stress pathways, thereby directly connecting the I/R cellular environment to the signaling cascades governing SG assembly^38,39^. The kidney’s prominent role as an oxygen sensor renders its tubular epithelium particularly sensitive to fluctuations in the intracellular redox environment, making it an especially relevant system in which to study stress-granule-mediated adaptive responses^36^. Despite growing recognition that SG formation occurs in renal proximal tubular cells during metabolic and ischemic stress and that modulation of SG dynamics may represent a novel therapeutic strategy for AKI^40^, the molecular factors governing SG regulation within kidney tissue remain poorly characterized.

Here, we show that SSB1 is selectively upregulated and recruited to G3BP1-marked SGs in a stress-dependent manner, where it functions as a negative modulator of SG dynamics under sublethal conditions. Loss of SSB1 enhances cellular sensitivity to oxidative stress without impairing proliferation following mild acute stress. We further identify stress-induced, G3BP1-dependent nuclear export of SSB1, highlighting coordinated regulation of its subcellular trafficking. Extending these cellular findings *in vivo*, we demonstrate marked induction and cytoplasmic granulation of SSB1 in the kidney following ischemia-reperfusion injury in mice. Collectively, these results establish SSB1 as an active regulator of cytoplasmic stress responses at both cellular and organismal levels, with potential implications for selectively targeting stress-adaptive pathways in cancerous, degenerative, and metabolic diseases.

## RESULTS

### Generation and validation of human RPE-1 knockout cell lines

We used a hTERT-immortalized human RPE-1 cell line to generate non-cancerous human knockout (KO) models for SSB1 (NABP2) and G3BP1. KO generation in RPE-1 cells has been well established, and these cells are also well suited for microscopic visualization of intracellular (nuclear and cytoplasmic) processes^41^. Using a CRISPR-Cas9-based gene-editing technique, we aimed to introduce frameshift mutations in both alleles of each target gene. By applying a series of gRNA constructs for each gene, we successfully created KO cell lines for SSB1 and G3BP1. Genotypic changes were verified by T7 endonuclease I-based assays and Sanger sequencing, whereas the absence of the target proteins was verified by Western blots and immunocytochemical visualization (**Fig. S1, Tables S1-S2**; see also Methods).

### SSB1 is embedded in cytoplasmic stress granules and negatively regulates stress granule formation

Our previous results indicated that SSB1 is included in cytoplasmic stress granules (SGs) in human cancerous (HeLa), tumorigenic (HEK-293T), and non-cancerous (HFF-1) cell lines upon acute oxidative stress induced by H_2_O_2_ treatment^25^. Here, we set out to define the contribution of SSB1 to SG formation using the above-described RPE-1 cell lines. First, we explored SG formation in wild-type (WT) RPE-1 cells upon treatment with various stressors (**Fig. 1**). Besides H_2_O_2_, we treated RPE-1 cells with sodium arsenite (NaAsO_2_) and potassium bromate (KBrO_3_), which are often used to induce direct oxidative stress^31,42^. Menadione (menadione sodium bisulfite) generates ROS through redox cycling while disrupting mitochondrial membrane potential and triggering cytochrome c redistribution to the cytosol, inducing indirect, internal oxidative stress^43^. DTT was used to induce endoplasmic reticulum stress by inhibiting protein folding through the reduction of disulfide bridges^44^. DTT is also responsible for ROS production by influencing cell signaling and the glutathione system^44^. Etoposide inhibits DNA topoisomerase II, leading to the accumulation of DNA breaks, where hSSB1 was shown to localize^4,7,45^. Thapsigargin irreversibly inhibits the SERCA Ca^2^□ pump, depleting endoplasmic reticulum calcium stores and triggering ER stress, unfolded protein response signaling, and often apoptosis^46^. We observed that SG formation became progressively more pronounced with increasing treatment times and doses of H_2_O_2_, arsenite, and DTT; whereas it remained negligible upon KBrO_3_, thapsigargin, and etoposide treatment (**Fig 1A-B**). The incorporation of SSB1 in SGs was evidenced by the significant enrichment of SSB1 signal in G3BP1-marked SGs upon treatment with various stressors (**Fig. 1C**). The signal of G3BP2, which was utilized to enable SG imaging independent of the presence of G3BP1^33^, was similarly enriched in G3BP1 SGs (**Fig. 1C, Fig. S2**). The strongest SG formation response was observed in menadione and arsenite (**Fig. 1A-B**); therefore, we used these stressors to define the contribution of SSB1 to SG formation by comparative analysis of WT, SSB1-KO, and G3BP1-KO RPE-1 cells (**Fig. 2**). We observed that SSB1-KO cells attained pronounced SG formation at a lower dose of menadione compared to WT cells (based on G3BP1 signal) and exhibited higher extents of SG formation at higher menadione doses than did WT cells (based on G3BP2 signal) (**Fig. 2A-B**). Moreover, we found that arsenite-induced SG formation, as monitored by G3BP2 signal, was enhanced in SSB1-KO cells compared to WT and G3BP1-KO cells (**Fig. 2D-E**). Collectively, these data define a negative SG regulator role for SSB1 under these conditions, in line with earlier implications regarding H_2_O_2_-induced SG formation in HeLa cells^25^.

**Figure 1.**
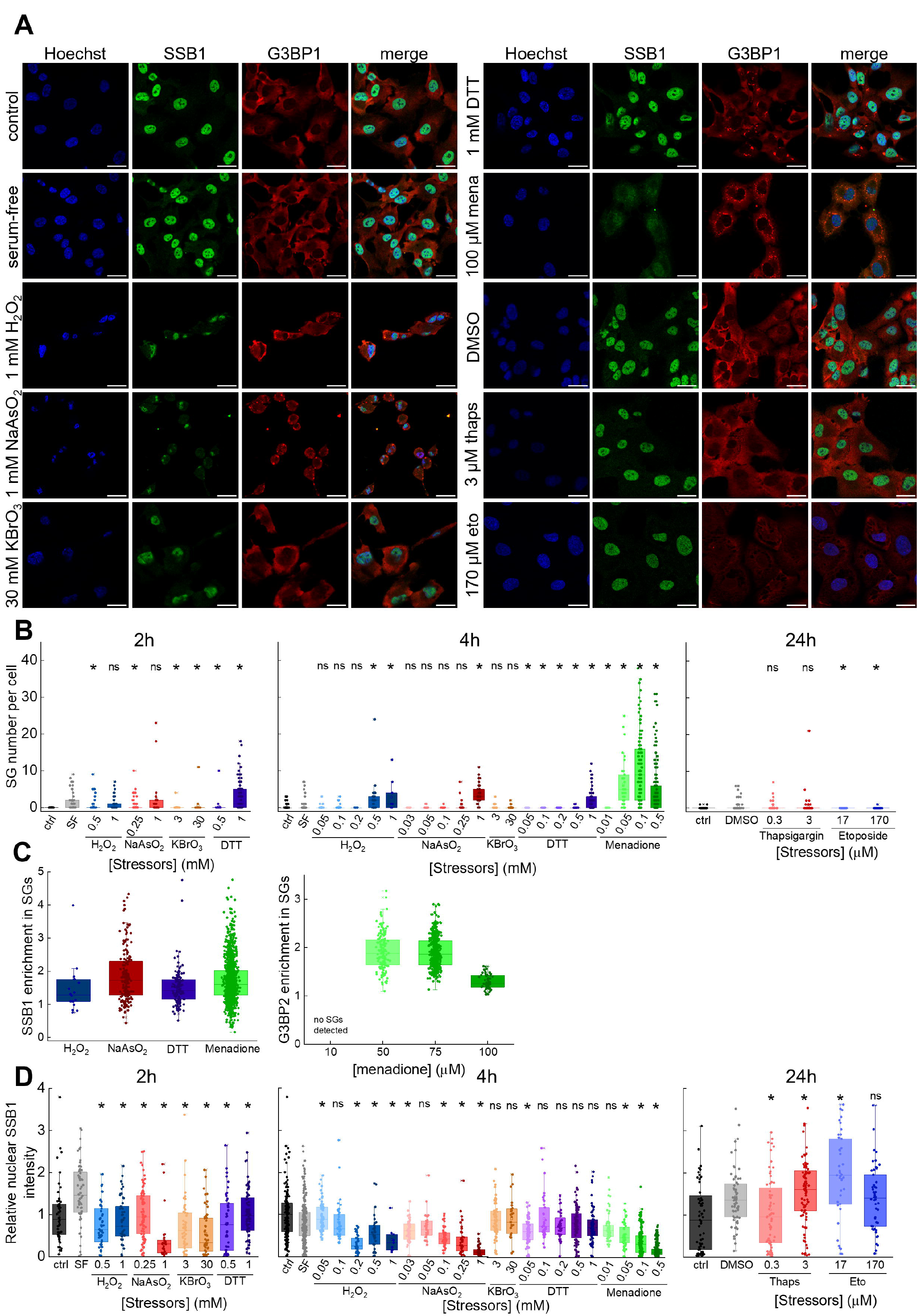
Cellular stress induces SSB1 enrichment in cytoplasmic stress granules and a decrease in nuclear SSB1 content. **(A)** Representative confocal immunocytochemical images of wild-type (WT) RPE-1 cells treated with various stressors. Blue channel shows nuclear Hoechst stain; green and red channels show endogenous SSB1 and G3BP1 stress granule (SG) markers, respectively. 4-h treatments are shown for serum-free control, H_2_O_2_, NaAsO_2_, KBrO_3_, DTT, and menadione (mena). 24-h treatments are shown for DMSO control, thapsigargin (thaps) and etoposide (eto). Scale bar: 30 μm. H_2_O_2_ and NaAsO_2_ treatment resulted in the appearance of rounded and morphologically compromised cells, consistent with reduced cell viability at these stressor concentrations. **(B)** Quantification of SG formation. Menadione induced the most robust SG formation of all stressors applied. Dots represent SG numbers from individual cells. Ctrl, untreated control; SF, serum-free treated controls. ‘*’ indicates significant difference from SF; ‘ns’, not significant (Mann-Whitney test, *p* < 0.05). **(C)** Enrichment of SSB1 (left) and G3BP2 (right) signals in SGs. Dots represent ratios of mean signal intensity inside individual stress granules detected *via* G3BP1 signal, compared to immediate surrounding cytoplasmic signal. In all cases, the population median was significantly higher than 1, indicating enrichment (one-sample Wilcoxon Signed Rank test, *p* < 0.05). For SSB1 enrichment analysis the following datasets were used: 1 mM H_2_O_2_, 1 mM NaAsO_2_, 1 mM DTT, 0.1 mM menadione; 4-h treatment time in all cases. Representative images of G3BP1 and G3BP2 co-staining are shown in **Fig. S2. (D)** Stress-induced changes in nuclear SSB1 intensity. Various stressors elicited a decrease in nuclear SSB1 intensity after 4-h treatment. Intensities were normalized to the means of untreated controls (ctrl). Dots indicate individual nuclear intensities. SF, serum-free treated controls. ‘*’ indicates significant difference from SF or DMSO controls; ‘ns’, not significant (Mann-Whitney test, *p* < 0.05).

**Figure 2.**
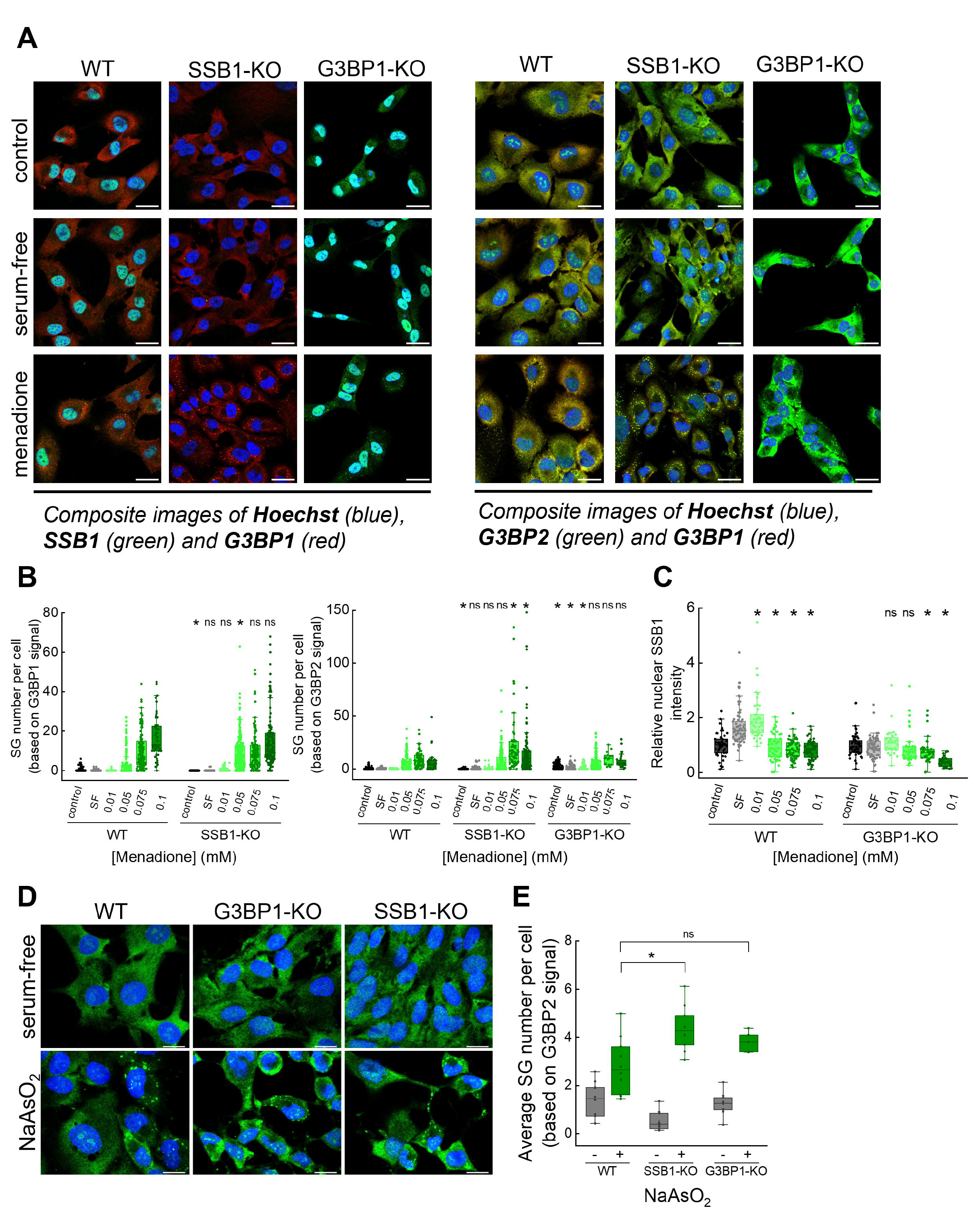
SSB1 negatively regulates stress granule formation. **(A)** Representative confocal immunocytochemical composite images of WT, SSB1-KO, and G3BP1-KO RPE-1 cells treated with 50 μM menadione for 4 hours. SF, serum-free treated control. Scale bar: 50 µm. **(B)** SSB1 negatively regulates the formation of menadione-induced stress granules. Dots represent SG numbers in individual cells. ‘*’ indicates significant difference between WT and KO cells at each condition; ‘ns’, not significant (Mann-Whitney test, *p* < 0.05). Menadione treatment was applied for 4 hours. **(C)** Menadione-induced changes in nuclear SSB1 intensity in WT and G3BP1-KO RPE-1 cells. Dots represent intensities in individual nuclei. ‘*’ indicates significant difference from SF; ‘ns’, not significant (Mann-Whitney test, *p* < 0.05). **(D)** Representative confocal immunocytochemical composite images of WT, SSB1-KO, and G3BP1-KO RPE-1 cells treated with 250 µM NaAsO_2_ for 2 h are shown, compared to serum-free treated controls. Blue signal shows nuclear Hoechst stain, green signal corresponds to G3BP2 staining. Scale bar: 10 µm. **(E)** SSB1 knockout enhances arsenite-induced stress granulation monitored through G3BP2 signal. Dots represent average SG numbers per cell from individual images. ‘*’ indicates significant difference from corresponding WT values; ‘ns’, not significant (Mann-Whitney test, *p* < 0.05).

### SSB1 shows stress- and G3BP1-dependent nuclear efflux

Our previous findings in HeLa cells demonstrated that H_2_O_2_ treatment triggers an initial rapid nuclear accumulation of SSB1, followed by a decrease in nuclear SSB1 signal on a longer (2 h) time scale, indicating possible nuclear efflux of the protein^25^. Consistently, the current study reveals that nuclear SSB1 levels in RPE-1 cells decrease in both a time- and dose-dependent manner following stressor exposure (**Figs. 1D, 2C**). Given the potential for functional crosstalk between SSB1 and G3BP1, we examined how G3BP1 influences the nucleo-cytoplasmic redistribution of SSB1. Intriguingly, we found that the nuclear SSB1 signal decrease, seen for WT RPE-1 cells upon both H_2_O_2_ and arsenite stress, was absent in G3BP1-KO cells in the case of H_2_O_2_ stress (**Fig. 3A-D**). Western blot-based detection of relative amounts of total cellular SSB1 confirmed no systematic change (or even a moderate increase upon short-time H_2_O_2_ treatment), confirming that the reduced nuclear SSB1 signal results from cytoplasmic protein accumulation rather than altered expression levels (**Fig. 3E-F**). Moreover, the lack of nuclear SSB1 intensity decrease upon thapsigargin and etoposide treatment was paralleled by a lack of SG formation under these conditions (**Fig. 1A-B, D**). Taken together, these findings demonstrate G3BP1-dependent cytoplasmic retention of SSB1 upon H_2_O_2_ stress, pointing to a functional interaction between these proteins and active involvement of SSB1 in cytoplasmic stress response.

**Figure 3.**
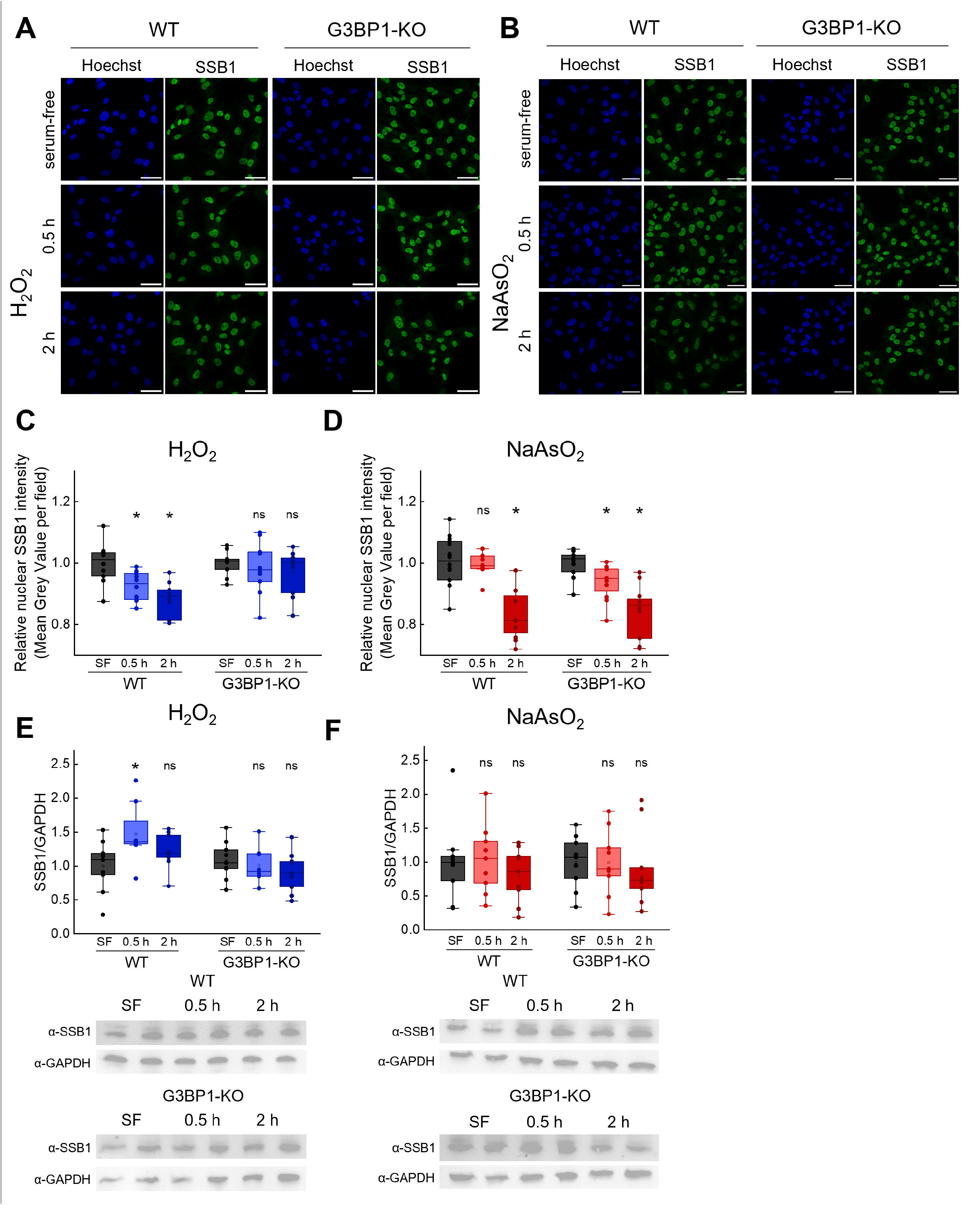
SSB1 undergoes G3BP1-dependent nuclear efflux upon hydrogen peroxide stress. **(A-B)** Representative confocal immunocytochemical images of WT and G3BP1-KO RPE-1 cells treated for 2 h with hydrogen peroxide (500 µM) and sodium arsenite (250 µM). Blue channel shows nuclear Hoechst stain, green channel shows endogenous SSB1. Scale bar: 50 µm. **(C-D)** Relative nuclear SSB1 intensities measured after 2-h hydrogen peroxide treatment. Data points represent mean intensity values measured from nuclei in individual fields. SF, serum-free treated controls. **(E-F)** Densitometric analysis of immunoblots monitoring SSB1 levels in WT or G3BP1-KO cells upon hydrogen peroxide and sodium arsenite treatment. Data points represent individual Western blot results. Representative immunoblots show two independent samples for each condition. All panels indicate results of two-sample *t*-tests. ‘*’ indicates significant difference from SF values (*p* < 0.05); ‘ns’, not significant.

### SSB1 and G3BP1 knockouts do not markedly affect acute stress survival and proliferation responses of the non-cancerous RPE-1 cell line

Next, we monitored the effects of SSB1 and G3BP1 knockouts on the acute stress response of RPE-1 cells by short-term exposure to different stressors (H_2_O_2_, arsenite, menadione), followed by monitoring cell viability using PrestoBlue assay performed immediately after exposure (**Figs. 4A** and **S3, Table S3**). We found that after surpassing a critical menadione concentration threshold, cell number declines markedly more abruptly in case of both SSB1-KO and G3BP1-KO cells, whereas WT cells exhibit a more gradual loss of viability (**Fig. 4A**). At the population level, this may represent a substantial difference, as WT cells are more likely to persist beyond the threshold, while the other populations undergo rapid cell death. Furthermore, SSB1-KO cells were significantly more sensitive to H_2_O_2_ compared to WT cells, but in the other applied conditions acute stress survival was unaffected by the knockouts (**Fig. 4A, Table S3**). Moreover, we monitored cell proliferation during a 6-day period after stressor treatment and washout (**Fig. 4B, Table S3**). G3BP1-KO cells showed generally slower growth compared to WT cells, but no marked knockout-dependent differences were apparent in the acute stress-induced proliferation responses of RPE-1 cell lines. Taken together, these results suggest that the pronounced developmental defects previously detected upon SSB1 ablation^10–15^ in animal models mostly result from chronic malfunctions in stress response, genome repair, and/or protein quality control.

**Figure 4.**
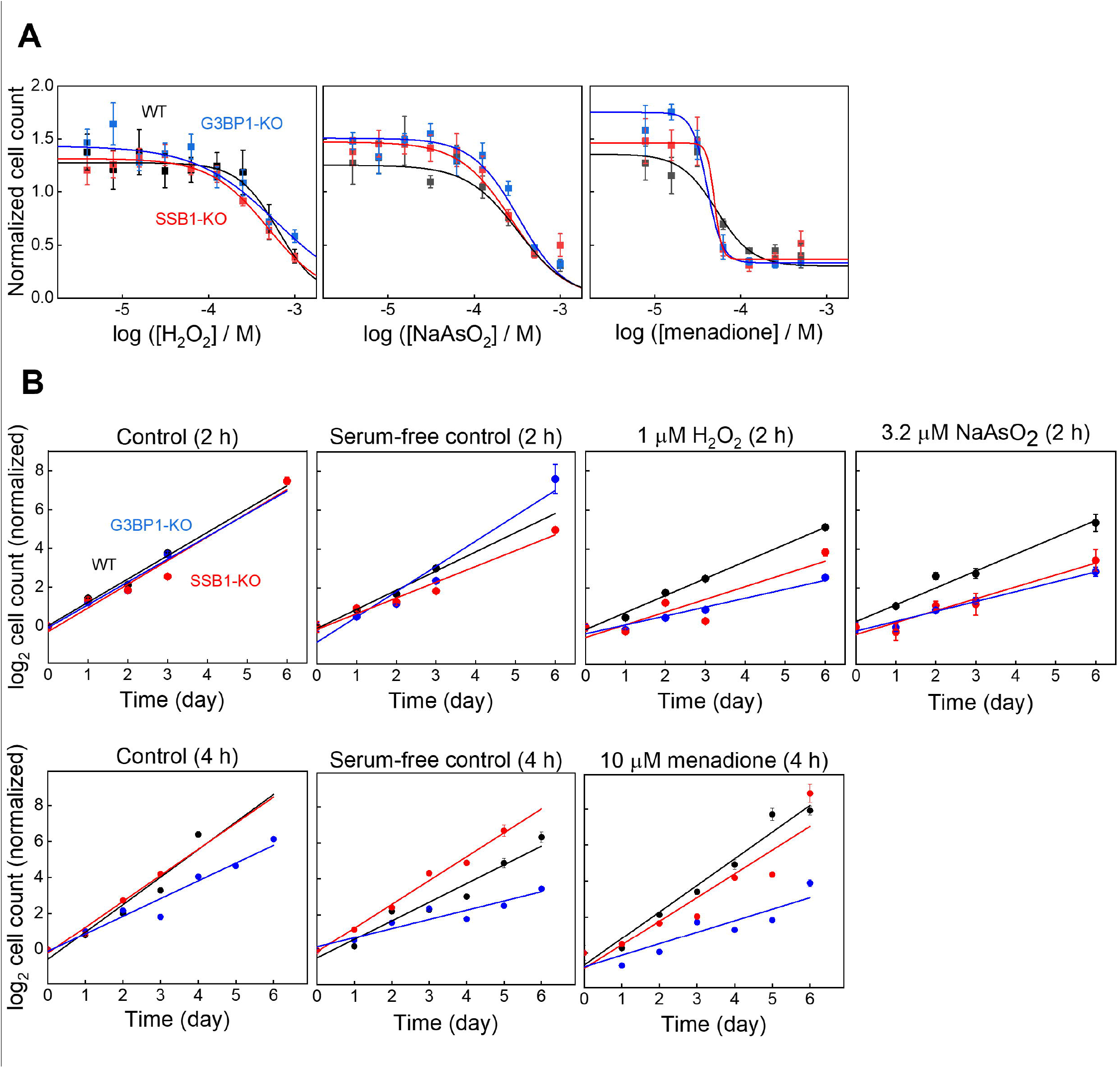
Cell survival and proliferation responses to acute stress are largely unaffected by SSB1 and G3BP1 knockouts. **(A)** PrestoBlue cell viability assays of WT and KO cells measured immediately after 2-h exposure to indicated stressors. Means ± SEM are shown for *n* ≥ 6 independent measurements. Data are normalized to serum-free controls (not shown). Solid lines show fits based on a standard dose-response function (*cf*. Materials and Methods). Best-fit log*LD50* values are listed in **Table S3. (B)** Proliferation of WT and KO cells monitored using PrestoBlue during a 6-day interval following stressor exposure and washout. Means ± SEM are shown for *n* ≥ 6 independent measurements. Data are normalized to 0-day values measured immediately after stress treatment. In the control groups, the medium was replaced with fresh complete medium for the indicated durations prior to measurement. Solid lines show linear fits. Best-fit proliferation rates are listed in **Table S3**. For all panels, cell counts were determined based on **Fig S3**.

### SSB1 undergoes intense cytoplasmic granulation and redistribution upon ischemia-reperfusion stress

SGs have been shown to be formed upon ischemia-reperfusion (I/R), indicating their functional contribution to this complex form of stress response^38,40^. Therefore, we assessed whether the cytoplasmic granulation of SSB1, detected in our earlier study^25^ and functionally analyzed in the current work in cell culture, also occurs in live-animal stress models. To this end, we applied renal I/R in mice and rats according to established protocols that mimic acute kidney injury (AKI)^47–54^. We found that SSB1 and G3BP1 protein levels significantly increased upon I/R stress in rat (SSB1) or both rat and mouse (G3BP1) kidney samples (**Fig. 5A**). However, the levels of SSB2, the SSB1 paralog used as a negative control, did not increase, highlighting the specific roles of SSB1, alongside those of G3BP1, in I/R response (**Fig. 5A**). Immunohistochemical investigation showed that the intensity of SSB1 increased upon I/R stress in mouse kidney samples (**Fig. 5B-C**). Importantly, in line with the cell culture observations, we detected cytoplasmic granulation of SSB1, which greatly increased upon I/R stress (**Fig. 5D**). Moreover, we assessed the subcellular distribution of SSB1 within renal epithelial cells and found that the quasi-homogenous distribution of SSB1-containing granules in unstressed animals along the apicobasal axis was markedly shifted toward the apical region in I/R animals (**Fig. 5E-F**). Collectively, these results constitute the first demonstration of SSB1’s involvement in cytoplasmic response to complex organismal stress, with corresponding stress-induced intracellular redistribution of the protein.

**Figure 5.**
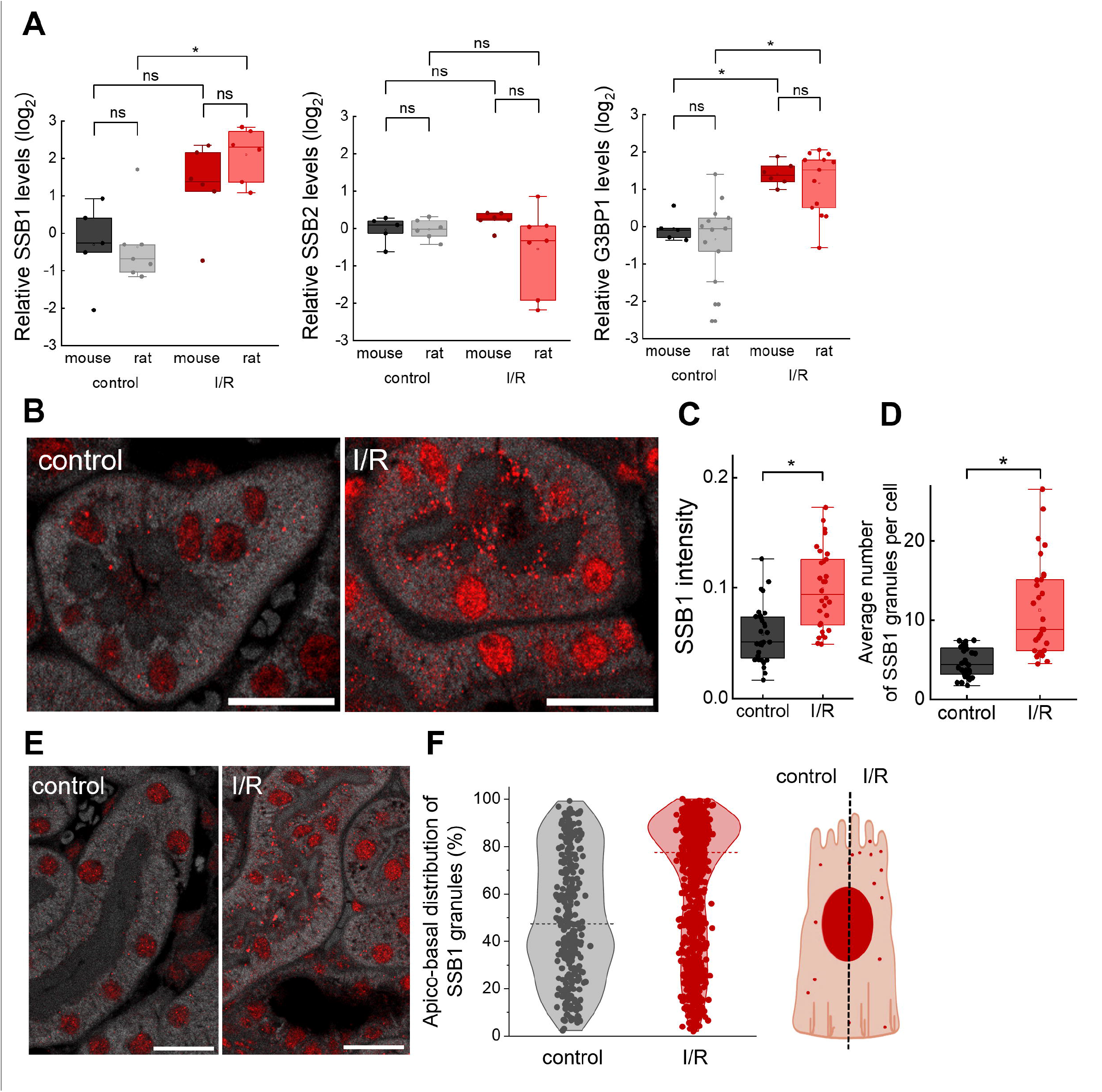
Renal ischemia-reperfusion stress induces elevation of SSB1 levels and massive cytoplasmic SSB1 granulation of SSB1 in rodents. **(A)** SSB1 and G3BP1 levels increase in rat and mouse kidney samples, whereas SSB2 levels are unchanged upon ischemia-reperfusion (I/R) (Western blot data; representative blots shown in **Fig. S4**). Data points represent samples from individual animals. **(B)** Representative immunohistochemical images of mouse kidney sections. Red stain indicates SSB1. Tissue morphology is visualized *via* paraformaldehyde autofluorescence captured in a separate channel. Scale bar: 20 µm. **(C)** Overall SSB1 intensity increase upon I/R in mouse kidney samples. Data points represent intensity data of individual images from 3 control and 3 I/R animals. **(D)** Average number of cytoplasmic SSB1 granules per cell in mouse kidney samples. Data points represent average granule numbers per cell from individual images of 3 control and 3 I/R animals. **(E)** Representative immunohistochemical images of mouse renal epithelial cells. Red stain indicates SSB1. Tissue morphology was visualized *via* paraformaldehyde autofluorescence captured in a separate channel. Scale bar: 20 µm. **(F)** Apico-basal distribution of SSB1 granules in renal epithelial cells (data from 3 control and 3 I/R animals). Each dot represents the relative position of an individual granule along the apico-basal axis. The cartoon depicts increased granulation in the apical region following I/R. In all panels, ‘*’ indicates significant difference from control values; ‘ns’, not significant (Mann-Whitney test, *p* < 0.05).

## DISCUSSION

Mammalian SSB1 has long been regarded as a nuclear protein with established key roles in genome maintenance^1–8,19^. However, we recently detected a significant decrease in nuclear SSB1 intensity (after a transient increase) in HeLa cells upon H_2_O_2_, arsenite, DTT, and menadione treatment, and the H_2_O_2_-induced decrease was also observed in HEK293T and HFF-1 cells^25^. In the current study we provide evidence that these changes indeed originate from stressor-induced efflux of SSB1 from the nucleus (as opposed to a decrease in cellular SSB1 levels), with SSB1’s cytoplasmic retention being selectively dependent upon G3BP1, a key SG constituent, upon H_2_O_2_ treatment (**Fig. 3**). These findings build on previous indications from a co-fractionation/mass spectrometry study reporting physical interaction between SSB1 and G3BP1^55^, and on our earlier report on SSB1 colocalization with G3BP1 SGs upon oxidative stress in HeLa, HEK, and HFF-1 cells^25^. Accordingly, in the current study we detected cytoplasmic G3BP1-SSB1 colocalization also in RPE-1 cells upon various forms of stress (H_2_O_2_, arsenite, DTT, menadione) (**Fig. 1**). Broadening the context of these findings, SSB1 interactors, traditionally implicated in nuclear genome maintenance, have also been detected in cytoplasmic SGs. Bloom syndrome helicase (BLM) has been identified as a negative regulator of SG formation^56^. In addition, a key component of the base excision repair pathway, hOGG1 (human 8-oxoguanine DNA glycosylase 1), has been shown to accumulate in SGs upon cadmium-induced oxidative stress^57^. Together, these observations suggest that SSB1-partner protein interactions beyond the SSB1-G3BP1 interplay may contribute to regulation of SG assembly, potentially involving hOGG1 and/or BLM recruitment to SGs by SSB1. Consistent with this proposition, we found that, alongside G3BP1, both BLM and hOGG1 are selectively enriched in SSB1 condensates^25^.

In conjunction with SSB1’s stress-induced efflux and incorporation into SGs, our current findings also define a negative regulator role for SSB1 in SG dynamics upon menadione and arsenite treatment in RPE-1 cells **(Fig. 2)**, building on our earlier indications for HeLa cells under oxidative stress^25^. We also show that SG formation and SSB1-linked regulation are stressor dose dependent and most pronounced in sublethal stress regimes **(Figs. 1B, 3-4, Table S3**). In line with this observation, a marked increase in SG numbers in the current study occurred at around the mentioned H_2_O_2_, arsenite, and menadione concentrations (**Fig. 1B**).

Notably, despite SSB1’s central roles in genome maintenance and ontogenesis, its ablation by siRNA knockdown in cultured U2OS and HeLa cells has been reported to result in detectable but modest sensitization to H_2_O_2_ and ionizing radiation^2,6,22,23,42^. In the current study, we also detected a significant but moderate sensitization for H_2_O_2_ in SSB1-KO RPE-1 cells compared to WT cells, while no effect of the knockout was seen on arsenite and menadione sensitivity (**Fig. 4, Table S3**). Intriguingly, in an earlier study, re-introducing the functionally partially compromised SSB1-C41S variant into SSB1-knockdown HeLa cells rescued WT sensitivity for gamma irradiation, but not for H□O□ treatment^21^, potentially pointing to effects from more extensive cytoplasmic damage in the latter case. Notably, in the current study we monitored cell survival colorimetrically immediately following acute stress, while earlier studies applied colony assays that require post-stress cell growth for detection. Nevertheless, for low stressor doses we did not detect SSB1 KO-induced differences upon cell proliferation following acute stress, suggesting that the effects of SSB1 ablation become pronounced in more chronic settings (**Fig. 4**).

In line with the above proposition, SSB1’s roles in SG dynamics and its upregulation in various cancerous tissues *in vivo*^25^ together indicate the utility of SSB1’s targeting for selective stress sensitization of cancer cells. Exposure to H_2_O_2_ (250 µM) caused 2-fold and 10-fold increases in SSB1 levels in HeLa^21^ and U2OS cells^2^, respectively, whereas an about 4 times elevation in SSB1 levels was seen in X-ray exposed humans (*i*.*e*., under genotoxic conditions)^58^. However, in the current study on non-cancerous RPE-1 cells we observed only a 1.5-fold increase in SSB1 levels after 0.5 h H_2_O_2_ treatment (at 500 µM stressor), which returned to base level at 2 h, and no increase was seen upon NaAsO_2_ stress (**Fig. 3E-F**). Moreover, moderate extents of SG formation were observed upon H_2_O_2_ treatment of cancerous HeLa and tumorigenic HEK-293T cells^25^, whereas very robust and prolific SG formation was seen for non-cancerous HFF-1 cells upon H_2_O_2_ stress^25^ and non-cancerous RPE-1 cells upon menadione treatment **(Fig. 1A-B)**. Taken together, these findings support an emerging view of SSB1 as a negative SG regulator that is overexpressed in transformed (malignant) cell lines, warranting more extensive studies to explore its utility in future therapeutic modalities.

Besides defining SSB1 as a regulator of SG dynamics that is characteristically redistributed within cultured cells upon stress, in the current study we also provide evidence for the same process occurring in kidney ischemia-reperfusion (I/R) injury, a complex organismal stress model (**Fig. 5**). As discussed below, the model we applied also reconstitutes features of acute kidney injury (AKI), a severe medical condition with unknown mechanism and missing therapy^34,35,37,39,59–61^. Besides oxidative stress, I/R typically creates conditions also involving endoplasmic reticulum (ER) stress, as also reported for AKI^38,62^. The prominent role of the kidney as an O_2_ sensor also makes this system well suited for probing sensitivity to the intracellular redox environment. Albeit SG formation is highly responsive to oxidative stress signals, and modulation of SG biogenesis and dynamics has been implicated as a novel approach to alleviate consequences of renal injuries, SG dynamics in the kidney are scarcely explored^40^. Together with SSB1’s association with G3BP1 stress granules^25,55^ (**Figs. 1-2**), our findings on SSB1’s involvement in renal I/R response lend further support to this concept.

eIF2α phosphorylation, a signal for SG assembly, has been observed to increase transiently in both the renal cortex and medulla upon reperfusion, peaking at 10 minutes and declining by 90 minutes after the onset of reperfusion^38^. In our study, we employed a 24-hour reperfusion model to better approximate a physiologically relevant AKI setting. Our detection of SSB1 granules in these conditions points to the protein’s sustained involvement in stress response, also consistent with the persistence of chronic ER stress following I/R, during which SGs could be maintained via PERK (PRKR-like ER kinase)-mediated phosphorylation of eIF2α^62^. Further evidence for the involvement of SGs in the stress response of renal epithelial cells comes from the finding that SGs were disrupted in G3BP1 siRNA-treated cultured proximal tubule cells upon azide or cisplatin treatment, accompanied by increased—but reversible—cell death associated with caspase-3 activation^40^. In rat proximal tubular cells, SG formation was observed following acute (0.5–3 h) treatment with glycolysis or mitochondrial inhibitors, whereas SGs showed a tendency to dissolve after 1–3 h of washout of mitochondrial inhibitors^40^. It will be important to determine in future studies whether early SGs reflect an adaptive response to reperfusion, whereas persistent SGs could potentially signal harmful consequences.

Perinuclear clustering of SGs is a common early response in stressed cells, promoting efficient mRNA sequestration and translational repression to conserve energy during acute injury^63^. Intriguingly, we observed SSB1 granules to concentrate in the apical region of kidney epithelial cells upon I/R (**Fig. 5E-F)**. eIF3h-labeled histological images taken shortly (10 min) after reperfusion showed perinuclear rather than apical localization of SGs in kidney epithelial cells^40^. Importantly, however, the time evolution of SG subcellular patterns has not been addressed in kidney I/R studies. Thus, it remains to be determined whether SGs redistribute to apical regions in longer reperfusion time regimes. Apical SG relocation, associated with restoration of epithelial cell polarity after stress, may enable localized translation of sequestered mRNAs encoding apical-specific proteins (*e*.*g*., those associated with brush border regeneration, transporter functions, or tight junctions), aiding epithelial barrier restoration^63^. Taken together, our current study defines cytoplasmic functions for SSB1 linked to both physiological stress response and severe medical conditions including cancerous transformation and tissue damage.

## MATERIALS AND METHODS

### Generation and validation of SSB1 (NABP2) and G3BP1 knockout (KO) human RPE-1 cell lines

sgRNA sequences for CRISPR-Cas9 genome editing were designed using Crispor and Chop-chop sgRNA prediction tools for NABP2, and the Brunello library for G3BP1^64^. Target sites were selected based on efficiency scores and localization of exons. sgRNA and primer sequences were verified using UCSC Genome Browser BLAST to prevent off-target annealing. Experimentally validated sgRNA sequences are shown in **Table S1**. Primers used for genotyping were designed using NCBI Primer Blast. Bbs1 restriction enzyme recognition sites were appended to oligonucleotides encoding sgRNA sequences to aid plasmid construction (see below). Oligonucleotides used for T7 endonuclease I (T7E1) assays (see below) and genotyping are listed in **Table S2**. Oligonucleotides were purchased from IDT. sgRNA-coding DNA sequences were inserted into the Cas9pX458-eGFP plasmid^65^ harboring an ampicillin resistance gene, the coding sequences for Cas9 and EGFP (both driven by a CMV promoter), a U6 promoter driving sgRNA expression, and two closely positioned BbsI restriction sites aiding sgRNA insertion. The generated sgRNA-coding plasmid constructs were purified using ZymoPURE Plasmid Miniprep Kit (D4209, centrifugation option) and verified by DNA sequencing.

5*10^5^ hTERT-immortalized human RPE-1 (retinal pigment epithelial) cells per well (in 6-well plates) were prepared for nucleofection using 500 µg of plasmid per well, using P3 Primary Cell Reagent and Supplementary Reagent (Lonza). Mixtures were then transferred into a nucleofection cuvette and transfected using a 4D-Nucleofector instrument (Lonza). 24 hours post-transfection, EGFP-positive cells were sorted using an Aria III FACS instrument. Individual positive cells were seeded onto 96-well plates and incubated until reaching confluency (about two weeks). Clonal populations were used for knockout validation.

PCR amplification of target sequences for T7E1 digestion-based screening^66^ was performed using cell lysates for template, along with primers listed in **Table S2**. PCR products, as well as T7E1 digestion products, were assessed in 1.5% agarose-TMB gels. Besides T7E1 assays and Sanger sequencing, cell lines were further validated using Western blot and immunocytochemistry (**Fig. S1**).

### Cell culture and treatments

RPE-1 cells were cultured in DMEM/F-12 medium (Sigma-Aldrich D6421) complemented with 10% FBS (Capricorn FBS-12A), 1% penicillin-streptomycin (Capricorn PS-b), and 0.01% hygromycin (Invivogen Hygromicin B-Gold). TrypLE Express was used to dissociate adherent cells for passage. For long-term storage, cells were frozen at -80°C in cell culture medium supplemented with 10% DMSO and then transferred to liquid nitrogen. Unless otherwise specified, stressors were applied in FBS-free medium to avoid the antioxidant effect of serum. Serum-free treatment controls were applied in the absence of stressors.

### Western blot

RPE-1 cells were lysed in Laemmli buffer (Bio-Rad) supplemented with 10 mM DTT. SDS-PAGE of cell lysates was performed using Mini-Protean Precast 4-20% gels (Bio-Rad 4561094). Primary antibodies used were α-human G3BP1 (from mouse, Abcam ab56574), α-human SSB1 (from rabbit, Sigma-Aldrich HPA044615), and α-human GAPDH (from rabbit, Sigma-Aldrich G9545). As secondary antibodies, anti-mouse-IgG HRP conjugate (Jackson 115-035-003) and anti-rabbit-IgG horseradish peroxidase (HRP) conjugate (Jackson 111-035-003) were used. Blots were stained using Immobilon Crescendo Western Blot HRP substrate (Sigma WBLUR0500). Bands of interest were analyzed using GelQuant Pro v12 by densitometry.

For rat and mouse kidney samples, total protein was extracted from frozen parts and from lyophilized homogenates of the same sample. Lysis buffer (1 M Tris, 0.5 M EGTA, 1% Triton X-100, 0.25 M NaF, 0.5 M phenylmethylsulphonyl fluoride, 0.5 M sodium orthovanadate, 5 mg/mL leupeptin, 1.7 mg/mL aprotinin, pH 7.4) was used to homogenize samples. Lysates were centrifuged at 13,000 rpm for 10 min at 4°C. Protein concentrations in supernatants were measured by detergent-compatible Bradford dye-binding method protein assay kit (Bio-Rad Hungary). 10 µg of denatured proteins were electrophoretically separated on 4-20% gradient Mini PROTEAN TGX SDS-polyacrylamide precast gels (Bio-Rad Hungary) and transferred to nitrocellulose membranes (TransBlot Turbo; Bio-Rad Hungary). Membranes were blocked in 5% w/v non-fat dried milk in Tris-buffered saline (TBS) for 1 hour at room temperature and probed with specific primary antibodies (α-human G3BP1 from mouse Abcam ab56574, α-human SSB1 from rabbit Sigma-Aldrich HPA044615, α-human SSB2 from rabbit Invitrogen PA5-100163) overnight at 4°C, followed by horseradish peroxidase-conjugated secondary antibodies (anti-mouse-IgG HRP conjugate Jackson 115-035-003, anti-rabbit-IgG HRP conjugate Jackson 111-035-003). Chemiluminescence detection was achieved using Molecular Imager VersaDoc MP 4000 System (Bio-Rad Laboratories, Hercules, CA, USA) using Luminata Forte (Millipore Corporation, Billerica, MA, USA) substrate. Bands were analyzed using GelQuant Pro v12 by densitometry.

### Immunocytochemistry and image analysis

Cells were fixed using 4% paraformaldehyde (dissolved in DPBS) for 20 minutes and permeabilized using 0.1% Triton X-100 in DPBS for 5 minutes at room temperature. Samples were then blocked using DPBS plus 5% BSA for 1 hour to reduce nonspecific binding. Cells were incubated with primary antibodies (identical to those used in Western blots, plus anti-human G3BP2 from rabbit, Proteintech 16276-1-AP) overnight at 4°C and then washed with DPBS. Secondary antibodies (anti-mouse-IgG Alexa488 conjugate (Invitrogen A11001) or anti-rabbit-IgG Alexa568 conjugate (Invitrogen A11011) were incubated with the cells for 1 hour, protected from light. Nuclei were stained using Hoechst 33342 (Thermo Scientific 62249, 1 mM) in DPBS for 15 minutes. Imaging was performed using a Zeiss LSM 710 confocal microscope (40x objective, NA 1.4) equipped with PMT detectors. A 405 nm / 30 mW diode laser, a 488 nm / 25 mW argon laser, and a 633 nm / 5 mW HeNe laser were used for excitation. Pinhole was set to 1 AU in all cases. SG number analysis was performed using scripts prepared in-house. Individual cells were detected by Cellpose based on G3BP1 signal^67^, and SGs were counted in each identified cell using the built-in thresholding algorithm of Fiji, also based on G3BP1 signal. Mann-Whitney analysis was performed on nuclear intensity and SG number data (significance level *p* < 0.05) using OriginPro 2018. For the analysis of SSB1 and G3BP2 enrichment in SGs, G3BP1 signal was used to detect individual SGs as ROIs using the built-in thresholding algorithm in Fiji. The mean signal intensity of SSB1 or G3BP2 was measured both inside the detected ROIs and in 0.4-μm bands surrounding them. Enrichment was calculated as a ratio of the two values. For the determination of nuclear SSB1 intensities, nuclei were detected as ROIs based on Hoechst signal. Integrated pixel density of SSB1 signal was measured for nuclear ROIs and normalized to indicated controls.

### Cell proliferation and stressor dose-dependent survival assays

For proliferation assays, 1,000 RPE-1 cells were seeded in wells of a 96-well plate in D-MEM/F-12 medium complemented with FBS, Penicillin-Streptomycin, and hygromycin, as described above for Cell culture and treatments. For stressor dose-dependent cell survival assays, 6,000 cells were seeded per well. Treatment was applied 24 hours after seeding. Stressors were applied in serum-free and antibiotics-free D-MEM/F-12 media. Stressor-free controls were set up in the same conditions. For proliferation assays, 10% diluted PrestoBlue reagent (in supplemented D-MEM/F-12 media) was added to the cells immediately after treatment (zero time point), or 24, 48, 72, and 144 hours after treatment. For dose-dependent survival assays, diluted PrestoBlue was added to the cells immediately after treatment. Cells were incubated with PrestoBlue for 2 hours unless otherwise specified. Fluorescence was measured at 590 nm with 560-nm excitation wavelength. All conditions were tested for at least *n* = 6 replicates.

Cell counts were calculated using a calibration curve measured independently from the proliferation and dose-dependent stressor survival analysis (**Fig. S3**). Cells were seeded in supplemented media in a two-fold dilution series (10 dilutions starting from 100,000 cells), then the media was replaced with 10% diluted PrestoBlue reagent after 24 hours. Fluorescent signal was measured after 1, 2, and 3 hours on independent samples (*n* = 6). Rectangular hyperbola (equation: *y* = *P*_1_\**x*/(*P*_2_ + *x*)) fitting was performed using *P*1 and *P*2 as floating parameters (best-fits obtained were *P*_1_ = 5693, *P*_2_ = 16858), where *y* is the PrestoBlue signal and *x* is the cell number. 2-hour PrestoBlue incubation time was applied in all analytical experiments. For dose-dependent survival analysis, a standard dose-response equation was used 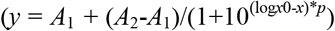 where *y* is the normalized cell count, *A*_1_ and *A*_2_ are bottom and top asymptotes, respectively, log*x*0 is log*LD*50 value, *x* is the stressor dose in log10 (M) scale, and *p* is Hill parameter) (**Fig. 4**).

### Rat and mouse renal ischemia-reperfusion

8-week-old male Charles River Wistar rats (Charles River Laboratories, Sulzfeld, Germany) and 6-week-old male C57BL/6J mice (Animalab Hungary Kft., Budapest, Hungary) were kept in groups of 3 (rats) or 6 (mice) in a temperature-controlled (24 ± 1°C) and humidity-controlled (relative 50 ± 10%) room under a 12h/12h light-dark cycle. Standard laboratory rodent chow and tap water were available *ad libitum*. General anesthesia was induced *i*.*p*. by ketamine (Richter Gedeon Plc., Budapest, Hungary; 75 and 90 mg/bwkg for rats and mice, respectively) and xylazine (Medicus Partner, Biatorbágy, Hungary; 10 mg/bwkg). During surgical procedures, body temperature was maintained at 37°C using a heating pad. Renal ischemia was accomplished by cross-clamping the left renal pedicle with an atraumatic vascular clamp for 45 and 25 min for rats and mice, respectively. Before the end of the ischemic time, contralateral nephrectomy was accomplished, followed by the removal of the clamp and 5 min observation of the left kidney to confirm complete reperfusion. 24 hours after reperfusion, animals were re-anesthetized and euthanized by exsanguination. Blood and kidney samples were collected. Sham-operated animals served as controls. Animal procedures were approved by the Committee on the Care of Laboratory Animals at Semmelweis University, Budapest, Hungary (PEI/001/1731-9-2015).

### Immunohistochemistry of mouse kidney samples

Kidney samples were rinsed in PBS and fixed using 4% PFA for at least 24 hours. Samples were then paraffin embedded and sectioned using a Leica rotation microtome (average thickness around 1 µm). Deparaffination was performed using xylene, while an ethanol dilution series (100%, 95%, and 70%) was used for rehydration following a published protocol^68^. Heat-induced antigen retrieval was performed in citrate buffer^68^ using a programable rice cooker, applying a 15-minute protocol three times, allowing the samples to cool slowly between cycles. 5% BSA – DPBS was used (1 h) for blocking and antibody dilution. The primary antibody (α-human SSB1, rabbit, Sigma-Aldrich HPA044615, 1:1000) was incubated overnight in a humid chamber, at 4°C. Alexa-488 labelled α-rabbit IgG was used (1:200, 2 hours) as a secondary antibody, then sections were washed 3 times in DPBS. Sections were mounted in Mowiol 4-88 and covered by 0.75 mm coverslips. Non-overlapping images from renal cortex were acquired by a Zeiss LSM 710 confocal microscope (40x Plan-Apochromat oil immersion objective, NA = 1.4). Samples had strong autofluorescence due to PFA fixation, which allowed us to capture tissue morphology in a separate channel.

### Quantification and statistical analysis

Microscopic images were processed using Fiji/ImageJ. Numerical data analysis and visualization was performed using OriginPro 2018. Pixel densitometry of electrophoretic gels and immunoblots was performed using GelQuant Pro software v12. Statistical analysis was performed in OriginPro 2018. Statistical tests and significance levels are indicated in figure legends.

## ACKNOWLEDGMENTS

We thank György Várady (HUN-REN RCNS) for assistance with FACS experiments, and Márton Nagyházi (HUN-REN RCNS) for the preparation of menadione sodium bisulfite. This work was supported by grants ELTE KMOP-4.2.1/B-10-2011-0002, and NKFIH ADVANCED 150087 to M.K. The project was supported by the NRDIO (VEKOP-2.3.3-15-2016-00007) grant to ELTE. J.P. was supported by the EKÖP-24 university excellence scholarship program (EKÖP-24-4-I-ELTE-184) of the Hungarian Ministry for Culture and Innovation from the source of the National Research, Development and Innovation Fund. J.P. was supported by the Cooperative Doctoral Program of the Ministry of Innovation and Technology financed from the National Research, Development and Innovation Fund. P.E. is a holder of the Bolyai Research Fellowship of the Hungarian Academy of Sciences (BO/00566/24). This work was completed in the framework of Project no. 2018-1.2.1-NKP-2018-00005 implemented with the support provided from the National Research, Development and Innovation Fund of Hungary, financed under the 2018-1.2.1-NKP funding scheme. Funded by the European Union HORIZON WIDERA 2023 IDP2Biomed grant agreement No. 101160233. Views and opinions expressed are however those of the author(s) only and do not necessarily reflect those of the European Union or the European Research Executive Agency (REA). Neither the European Union nor the granting authority can be held responsible for them. This research is carried out and financed within the framework of the second Swiss Contribution MAPS. This work was funded by the following grants to A.F.: project no. TKP2021-EGA-24 has been implemented with support from the Ministry of Innovation and Technology of Hungary from the National Research, Development and Innovation Fund, financed under the TKP2021-EGA (Thematic Excellence Programme 2021 – Health Subprogramme) funding scheme. The work was also supported by the STAGE 2024-1.2.3-HU-RIZONT-2024-00056 grant of the National Research, Development and Innovation Fund of Hungary and the MTA-SE “Lendület” Research Grant LP2021-3/2021 of the Hungarian Academy of Sciences. This project has received funding from the HUN-REN Hungarian Research Network. The authors declare no competing interests.

