## Supplementarty Information for "SSB1 links stress granule regulation to cellular stress responses and renal ischemia-reperfusion injury"

### SUPPLEMENTARY FIGURE LEGENDS

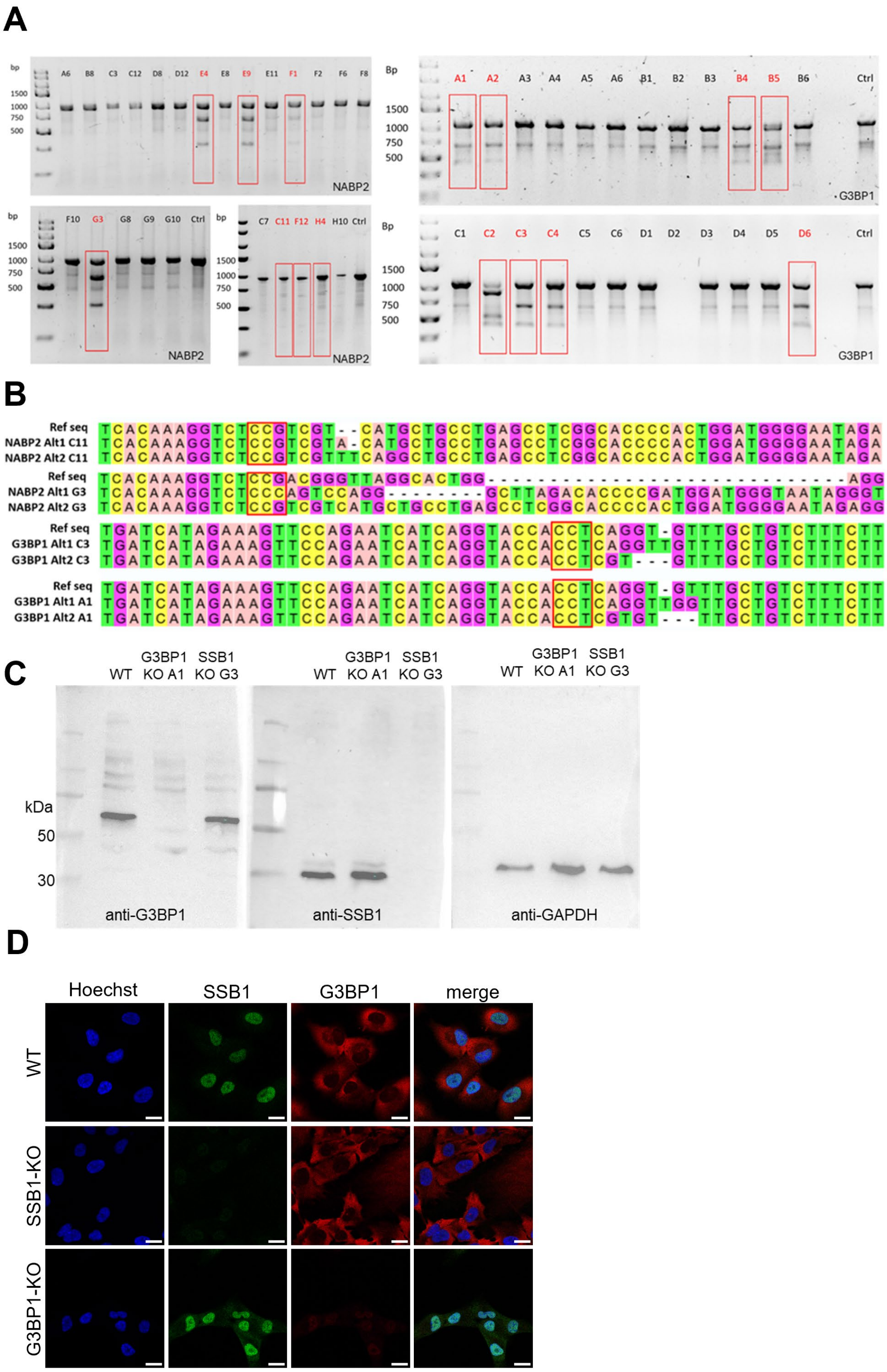


**Figure S1. Generation and validation of SSB1-KO (NABP2-KO) and G3BP1-KO RPE-1 cell lines.** **(A)** Representative T7 endonuclease I (T7E1) assay of single-cell clones resulting from NABP2 (exon 2 targeted) and G3BP1 (exon 4 targeted) knockout experiments. Positive results are shown in red boxes.

Single-band PCR product indicates that no mutation had occurred (unsuccessful knockout), whereas three bands indicate successful knockout as T7E1 cleaves the heteroduplex PCR product (mutated DNA) into two shorter fragments, visible below the original band. **(B)** Sanger sequencing results for compound heterozygote single-cell clones. Both alleles (Alt1 and Alt2) are null mutants, *cf.* reference sequences. The PAM sequence used for CRISPR-Cas9 editing is shown in red boxes. **(C)** Western blot validation of SSB1-KO (G3) and G3BP1-KO (A1) clones. **(D)** Representative immunocytochemical images of WT, SSB1-KO, and G3BP1-KO RPE-1 cell lines. Cells were stained in the absence of stress. Scale bar: 20 μm.

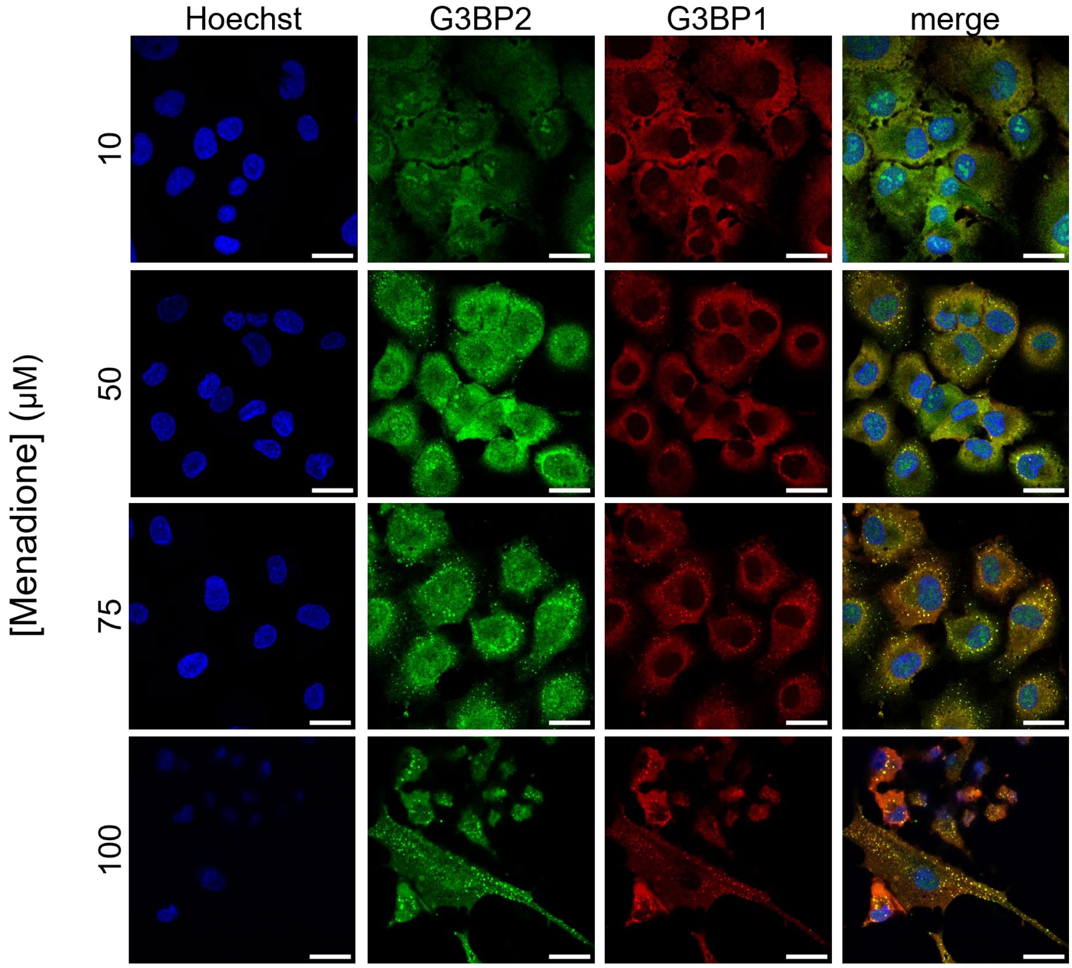


**Figure S2. Both G3BP1 and G3BP2 signals are suitable for visualization of stress granules.** Shown are representative confocal immunocytochemical images of WT RPE-1 cells treated with indicated concentrations of menadione for 4 h. Blue channel shows nuclear Hoechst stain; green and red channels show G3BP2 and G3BP1 stress granule (SG) markers, respectively. Scale bar: 30 µm.

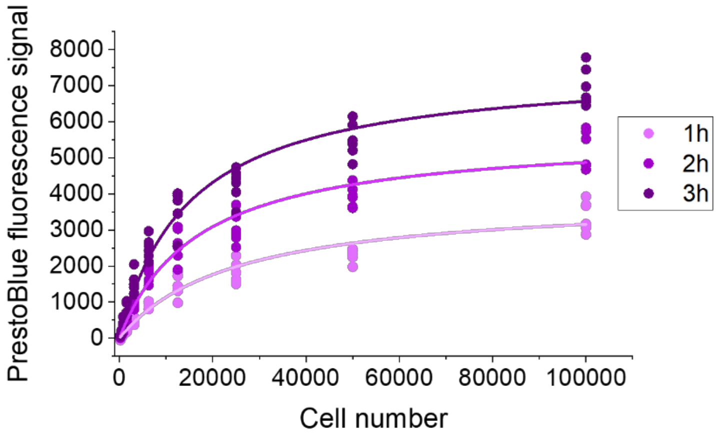


**Figure S3. PrestoBlue signal calibration.** PrestoBlue fluorescence signals at the indicated incubation times are shown as a function of input RPE-1 cell count. Calibration curves (solid lines) were obtained by applying rectangular hyperbolic fits to the data using the equation $y = P_1*x/(P_2+x)$ where $y$ is PrestoBlue signal, $x$ is input cell count, and $P_1$ and $P_2$ are floating parameters. Best-fit values obtained were $P_1 = 5693$ and $P_2 = 16858$.

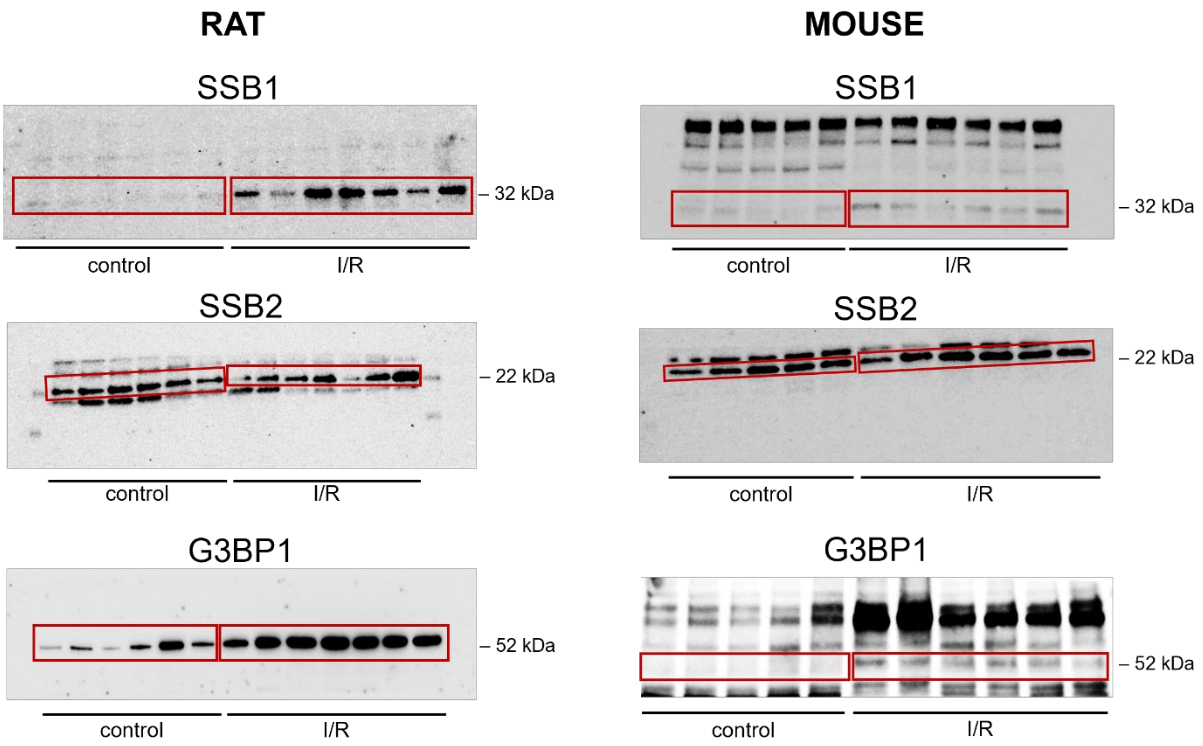


**Figure S4. Immunoblots of rat and mouse kidney samples from control and I/R animals.** Samples containing 10 µg total protein were loaded in each well. Lanes represent individual animals. Red boxes indicate bands corresponding to previously established apparent molecular weights of the analyzed proteins, as indicated beside the blots [1,2]. Results of band intensity analysis are shown in **Fig. 5A**. The appearance of aspecific bands in mouse G3BP1 immunoblots can be attributed to cross-reactivity of the applied anti-mouse secondary antibody with endogenous mouse immunoglobulins present in the tissue sample. In the case of mouse SSB1 immunoblots, the aspecific bands are of uncertain origin; however, the consistent direction and extent of I/R-induced changes in SSB1 amounts determined from Western blots (**Fig. 5A** and this figure) and immunohistochemical analysis of kidney epithelial cells (**Fig. 5C**), together with the similarity between subcellular (nuclear and cytoplasmic) distributions observed in stressed mouse kidney and cultured human cells (**Figs. 1, 2,** and **5**), demonstrate that the signals detected in histological sections indeed originate from labeling of SSB1 by the applied antibodies.

### SUPPLEMENTARY TABLES

**Table S1: Experimentally validated sgRNA sequences**
Asterisks indicate sgRNA sequences with successful targeting.

| **Target gene** | **Exon** | **Location** (human hg38 ref.) | **sgRNA** (presented as cDNA sequence, 5'-3') | **PAM sequence** |
|---|---|---|---|---|
| NABP2 | 2 | chr12: 56224844 -56224863 | AGGCTCAGGCAGCATGACGA* | CGG |
| NABP2 | 3 (#1) | chr12: 56225437-56225456 | CAGCATCAATATCTCTGTCT | GGG |
| NABP2 | 3 (#2) | chr12: 56225489 -56225508 | GACATTATCCGGCTCACCAA | AGG |
| NABP2 | 4 | chr12: 56225401-56225420 | TGAGGTTCGGACCTGCAAAG | TGG |
| G3BP1 | 4 | chr5: 151795507-151795526 | AGAAAGACAGCAAACACCTG* | AGG |
| G3BP1 | 8 | chr5: 151790965-151790984 | TACCACACCATCATTTAGCG | TGG |

**Table S2: Primers used for PCR amplification**
Asterisk indicates amplicon of aspecific targets.

| **Amplicon** | **Forward primer (5'-3')** | **Reverse primer (5'-3')** |
|---|---|---|
| NABP2 Exon 2 | CTCTGATCCCGATTGGCTGG | CTGAGTGGAAAGAGGCCCG |
| NABP2 Exon 3 (#1) | GACGACGGAGACCTTTGTGAA | CACCCTATGCTGCCTCCTTG |
| NABP2 Exon 3 (#2)* | CCAGTGGGGTGCCGAGGCTC | ACACCCTATGCTGCCTCCTTGC |
| NABP2 Exon 4 | CCAGTGGGGTGCCGAGGCTC | ACACCCTATGCTGCCTCCTTGC |
| G3BP1 Exon 4 | GTGGTAGAGCCTGCATTAGATAACTGCC | GGTCTCTGTCCACTTCATATTCCCTC |
| G3BP1 Exon 8 | GGGGGATTGGATTCAAATGGAAA | GGGCAAGTGGTGACAGCTGC |

**Table S3: Dose-dependent stressor survival and proliferation parameters of RPE-1 cell lines**.

| Analysis/stressor | Treatment time (h) | WT | SSB1-KO | G3BP1-KO |
|---|---|---|---|---|
| **log *LD*50 (M)** | | | | |
| $H_2O_2$ | 2 | -3.19 ± 0.04 | -3.31 ± 0.03* | -3.21 ± 0.09 |
| $NaAsO_2$ | 2 | -3.49 ± 0.09 | -3.57 ± 0.06 | -3.48 ± 0.05 |
| menadione | 4 | -4.28 ± 0.12 | -4.31 ± 0.03 | -4.39 ± 0.02 |
| **Proliferation slope (log 2 growth/day)** | | | | |
| Untreated control | 2 | 1.20 ± 0.12 | 1.22 ± 0.14 | 1.17 ± 0.17 |
| Untreated control | 4 | 1.53 ± 0.25 | 1.45 ± 0.14 | 0.98 ± 0.1* |
| Serum-free media | 2 | 0.99 ± 0.08 | 0.81 ± 0.08 | 1.30 ± 0.17 |
| Serum-free media | 4 | 1.04 ± 0.11 | 1.33 ± 0.07 | 0.52 ± 0.08* |
| 1 µM $H_2O_2$ | 2 | 0.87 ± 0.04 | 0.65 ± 0.18 | 0.46 ± 0.06* |
| 3.2 µM $NaAsO_2$ | 2 | 0.87 ± 0.09 | 0.61 ± 0.09 | 0.51 ± 0.05* |
| 10 µM menadione | 4 | 1.48 ± 0.12 | 1.32 ± 0.23 | 0.64 ± 0.14* |

Means ± SEM are shown for $n = 6$. Asterisks (*) indicate significant difference from WT values (two-tailed *t*-test, $p < 0.05$).
